# Mechanism of the ANT-mediated transport of fatty acid anions across the inner mitochondrial membrane

**DOI:** 10.1101/2022.06.27.497434

**Authors:** Jürgen Kreiter, Zlatko Brkljača, Sanja Škulj, Sarah Bardakji, Mario Vazdar, Elena E. Pohl

**Affiliations:** Institute of Physiology, Pathophysiology, and Biophysics, Department of Biomedical Sciences, University of Veterinary Medicine, Vienna, Austria; Division of Organic Chemistry and Biochemistry, Rudjer Bošković Institute, Zagreb, Croatia; Department of Chemistry, Faculty of Science, University of Zagreb, HR-10000 Zagreb, Croatia; Department of Mathematics, University of Chemistry and Technology, 166 28 Prague, Czech Republic

## Abstract

The additional protonophoric function of the mitochondrial adenine nucleotide translocase (ANT1) is now recognized. However, the molecular mechanism remains controversial. Fatty acid (FA) cycling hypothesis postulates that FAs transport protons across the inner mitochondrial membrane to the matrix by a flip-flop, whereas ANT1 facilitates the translocation of FA anions (FA^-^) back to the intermembrane space. By a combined approach involving measurements of current through the planar lipid bilayers reconstituted with recombinant ANT1, site-directed mutagenesis and molecular dynamics simulations, we show that FA^-^ is initially caught by R59 on the matrix side of ANT1, then moves along the positively charged protein-lipid interface, and binds to R79, where it is protonated in the hydrated cavity in the presence of D134. R79 is crucial for the competitive binding of ANT1 substrates (ATP and ADP) and inhibitors (carboxyatractyloside, bongkrekic acid). The binding sites are well-conserved in mitochondrial SLC25 members, implying a general transporting mechanism for FA anions.

## Introduction

The proton gradient Δμ(H^+^) across the inner mitochondrial membrane (IMM) is coupled with ATP production. The adenine nucleotide translocase (ANT, also AAC for ADP/ATP carrier) exchanges cytosolic ADP for matrix ATP to maintain energy production and supply. Dissipation of Δμ(H^+^) without the formation of ATP (uncoupling) is mediated by uncoupling protein 1 (UCP1), which produces heat during non-shivering thermogenesis in the brown adipose tissue of hibernating animals and newborns ^1^. Several proteins of the SLC25 superfamily, including UCP2, UCP3 ^2,3^ and ANT1 ^4-7^, transport protons although the exact biological function of the transport is unclear. The H^+^ turnover numbers of UCP1-UCP3 and ANT1 in the presence of arachidonic acid are similar, indicating a similar potency for H^+^ transport ^6,8-10^.

Several mechanistic models have been proposed to account for the protonophoric activity of UCPs and ANT in the presence of FA. In the FA cycling model (Fig. 1a), protons are transported by a flip-flop of the protonated fatty acids (FAH) ^11^, which quickly diffuse through the lipid bilayer ^12,13^. FA^-^ transport back to the intermembrane space is accelerated by protein in the FA cycling mechanism. In contrast, the H^+^ buffering model proposes that FA anions, bound inside the protein, provide an additional negative charge to accelerate H^+^ transport by proton wire ^14^. Finally, the FA shuttling model, which has only been proposed for UCP1 ^15^, holds that the FA is bound flexibly within a protein. Upon FA^-^ protonation on the cytosolic side, the FA swivels to the matrix side, releasing the proton and flipping back to bind a new H^+^.

**Figure 1.**
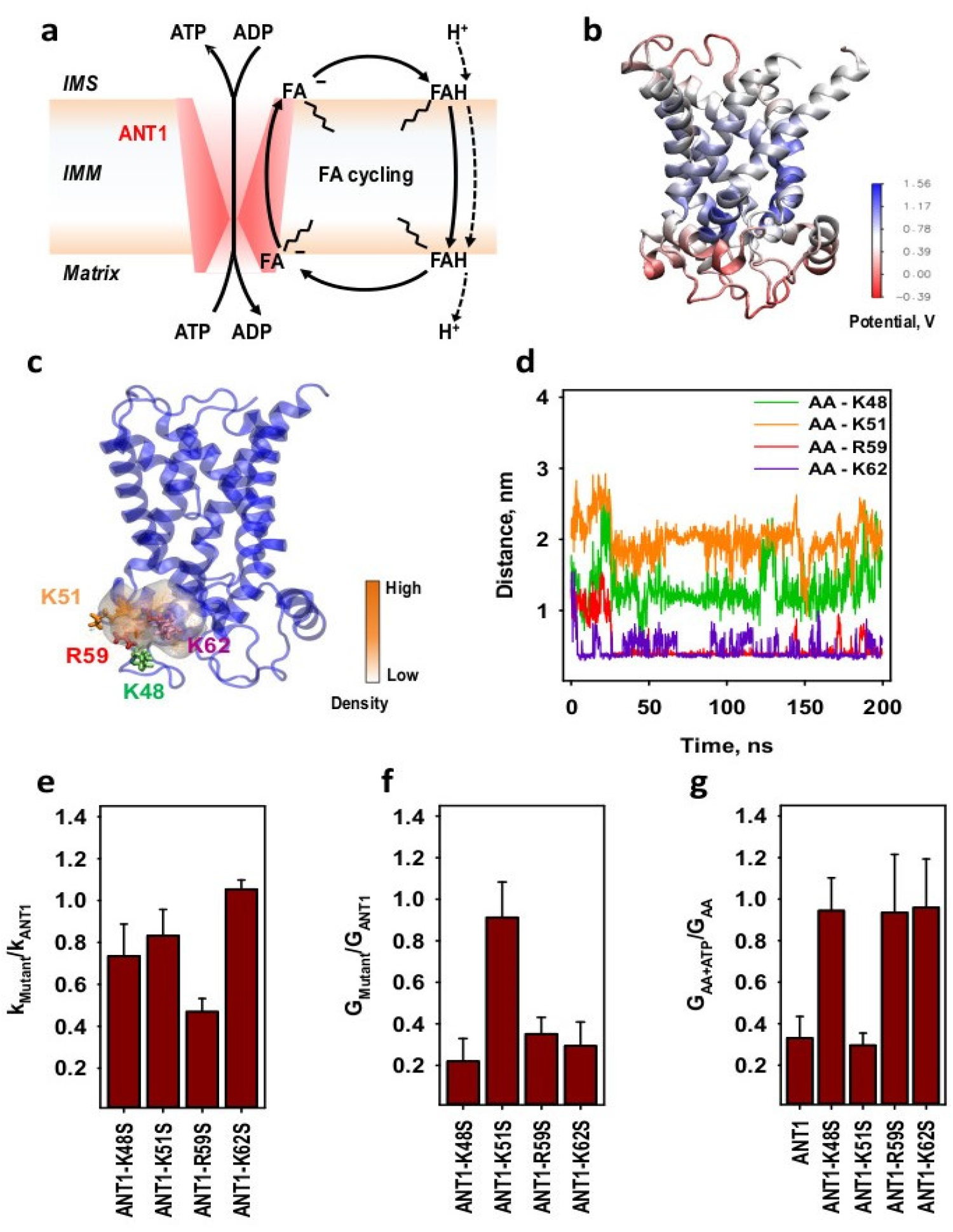
The fatty acid anion (FA^-^) binds to a set of arginine and lysine residues at the matrix side of ANT1. (a) The fatty acid cycling model, in which ANT1 is proposed to facilitate the transport of FA^-^ across the inner mitochondrial membrane (IMM) from the mitochondrial matrix to the intermembrane space (IMS). (b) The calculated electric surface potential of ANT1 (PDB:1OKC). (c) Number density map of the FA anion at the matrix side of ANT1 (PDB:1OKC) determined by non-equilibrium molecular dynamics simulations. Amino acids K48 (green), K51 (orange), R59 (red) and K62 (purple) are displayed in licorice. (d) Distances between the FA anion head and K48 (green), K51 (orange), R59 (red) and K62 (purple), extracted from the MD simulations in (c). (e) The ratio of the ADP/ATP exchange rate, k_Mutant,_ of the ANT1 mutants (ANT1-K48S, ANT1-K51S, ANT1-R59S and ANT1-K62S) relative to the exchange rate (k_ANT1_) of ANT1 measured with ^3^H-ATP. (f) The ratio of the total membrane conductance, G_Mutant_, of the membranes reconstituted with ANT1 mutants (ANT1-K48S, ANT1-K51S, ANT1-R59S and ANT1-K62S) to G_ANT1_ of the membranes reconstituted with ANT1 in the presence of arachidonic acid (AA) in the membrane. (g) The ratio of the total membrane conductance, G_AA+ATP_, in the presence of AA and ATP to the total membrane conductance, G_AA_, in the presence of AA only, calculated for the bilayer membranes reconstituted either with ANT1 or ANT1 mutants (ANT1-K48S, ANT1-K51S, ANT1-R59S and ANT1-K62S). In the experiments, the lipid concentration was 1.0 mg/ml (e) or 1.5 mg/ml (f, g). Protein concentration was 8.5 – 9 μg/(mg of lipid) (e) or 4 μg/(mg of lipid) (f, g). Membranes were made of PC:PE:CL (45:45:10 mol%) (e - g) and reconstituted with 15 mol% AA (f, g). Buffer solution contained 50 mM Na_2_SO_4_, 10 mM Tris, 10 mM MES and 0.6 mM EGTA at pH = 7.34. Temperature was 296 K (e) or 306 K (f, g). Data are the mean ± SD of at least three independent experiments.

FA cycling model also describes the activation and inhibition of FA-mediated H^+^ transport by ANT1 (Fig. 1a) ^6^, but the molecular details of the mechanism by which ANT1 transports FA^-^ back to the intermembrane space have remained obscure. We have now used the crystallographic structure of ANT1 ^16,17^ and a series of MD simulations ^18,19^ to design specific recombinant ANT1 mutants and analyze the transport of FA^-^. The work uncovers the molecular mechanism by which ANT1 transports FA^-^s. As the pathway utilizes highly conserved amino acids, it may also be used by uncoupling proteins to transport FA^-^.

## Results

### 1. Identification of the initial AA^-^ binding site on the ANT1 matrix side

Based on the surface potential analysis of ANT1 (Fig. 1b), we identified four positively charged lysine and arginine residues (K48, K51, R59, and K62) on the matrix side of ANT1 as most likely to represent an initial binding site for FA anions. Molecular dynamics simulations confirmed that the AA^-^ is trapped in the vicinity of these amino acids (Fig. 1c and Video 1). Fig. 1d shows that the average distances between the AA^-^ and K48 or R59, or K62 are small enough to ensure electrostatic interactions via salt bridge interactions, whereas the distance between AA and K51 is significantly higher. Thus, non-equilibrium MD simulation experiments indicate that K51 does not participate in AA^-^ trapping.

Using site-directed mutagenesis, we produced ANT1 mutants in which we substituted these amino acids by the polar but neutral serine (K48S, K51S, R59S, and K62S). The mutations had only slight effect on the ANT1-mediated ADP/ATP exchange rate apart from ANT1-R59S (Fig. 1e, Suppl. Fig. 1 and Suppl. Table 1), which is involved neither in binding the substrates nor in the matrix salt bridge network ^18,20^. Measurements of membrane current revealed that the total conductance of membranes reconstituted with the K48S, R59S, and K62S mutants in the presence of arachidonic acid (AA), G_Mutant_, decreased by 70 % but was almost not affected in K51 mutant (Fig. 1f and Suppl. Fig. 2). This confirms that K51 does not participate in the trapping of FA^-^. ATP has no inhibitory effect in most mutants (Fig. 1g and Suppl. Fig. 2), indicating the loss of coupling between ADP/ATP exchange and AA^-^ trapping.

### 2. AA^-^ slides along a positively charged surface at the protein-lipid interface

Previously we have shown that ANT1 has a considerable positive surface potential at the protein-lipid interface that is involved in binding AA^-^ on the matrix side ^6^. It extends deep into the centre of the bilayer membrane and includes the substrate-binding centre of the protein ^18^. To reveal the path for the AA^-^ translocation, we constructed density maps using a series of unbiased MD simulations in which the AA^-^ was randomly placed in the vicinity of ANT1 along the positively charged electrostatic potential. Figure 2a shows the number density of the AA^-^ at different positions in ANT1, which implies that AA^-^ may slide along the protein-lipid interface to the substrate-binding centre of ANT1, which includes at least three charged arginines (R79, R137 and R279) and two lysines (K22 and K93). Importantly, we performed the simulations in the presence of cardiolipin, an essential lipid in the inner mitochondrial membrane. This binds tightly to ANT1 and introduces negative charges to the lipid-protein interface ^21,22^.

**Figure 2.**
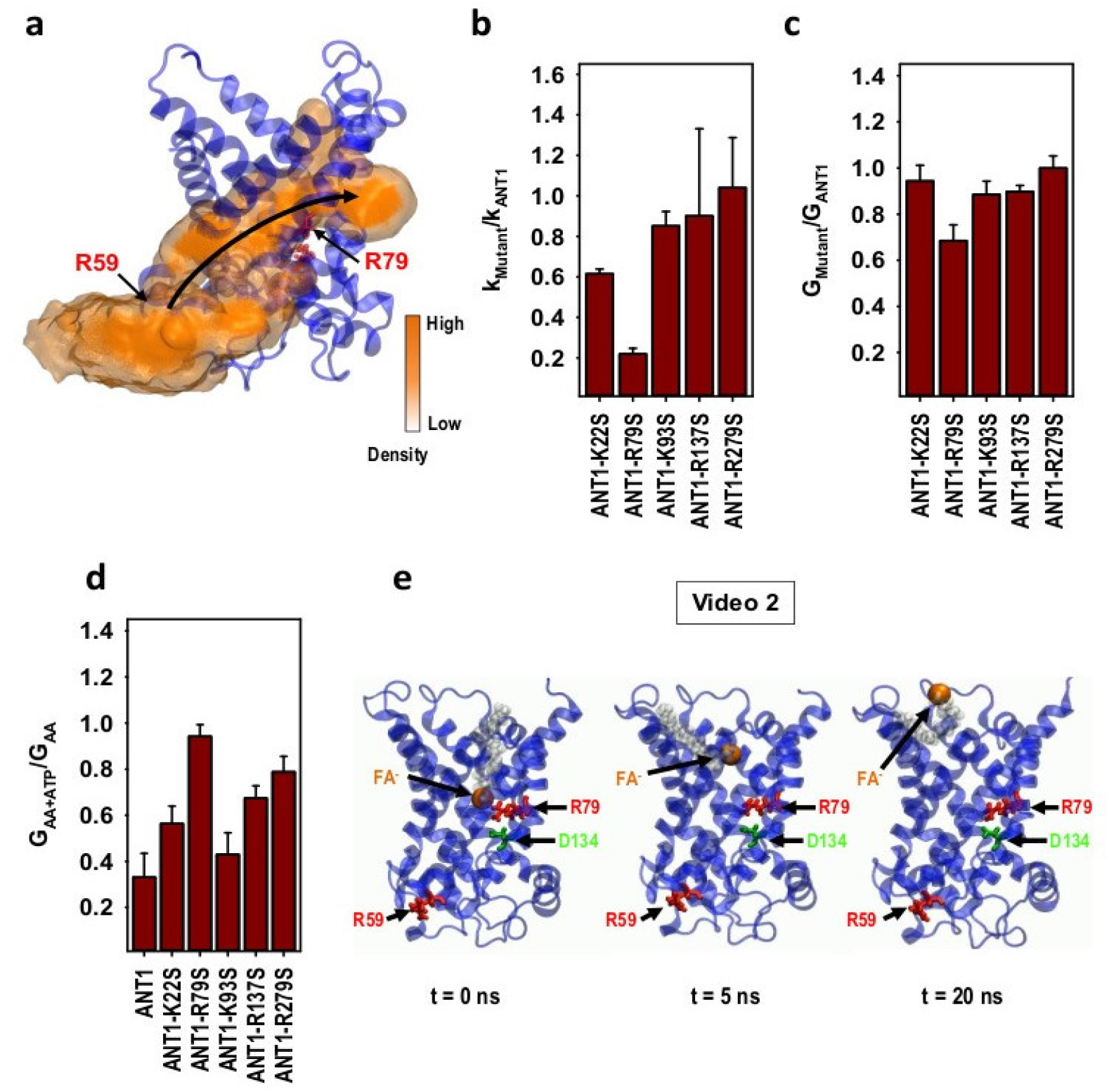
The fatty acid anion (FA^-^) slides along the protein-lipid interface. (a) Number density map of the FA anion at the surface of ANT1 (PDB:1OKC) determined by non-equilibrium molecular dynamic simulations. (b) The ratio of the ADP/ATP exchange rate, k_Mutant_, of the ANT1 mutants (ANT1-K22S, ANT1-R79S, ANT1-K93S, ANT1-R137S and ANT1-R279S) to the exchange rate, k_ANT1_, of ANT1 measured with ^3^H-ATP. (c) The ratio of the total membrane conductance, G_Mutant_, of the ANT1 mutants (ANT1-K22S, ANT1-R79S, ANT1-K93S, ANT1-R137S and ANT1-R279S) to the G_ANT1_ of the membranes reconstituted with ANT1, measured in the presence of AA in the membrane. (d) The ratio of the total membrane conductance, G_AA+ATP_, in the presence of AA and ATP to the G_AA_, in the presence of AA only, calculated for the bilayer membranes reconstituted either with ANT1 or ANT1 mutants (ANT1-K22S, ANT1-R79S, ANT1-K93S, ANT1-R137S and ANT1-R279S). (e) Series of snapshots of the FA anion placed in the vicinity of R79 inside the hydrated cavity from the cytosolic side. Snapshots are extracted from MD simulations shown in Video 1. Images were taken in the beginning (t = 0 s), after t = 5 ns and at the end of the simulation (t = 20 ns). The FA^-^ tail is displayed by van-der-Waals surfaces in grey, the FA^-^ head - as a ball in orange. Amino acids R59 (red), R79 (red) and D134 (green) are displayed in licorice in the structure of ANT1 (PDB:1OKC). Experimental conditions are the same as in Figure 1.

To test the next binding site at the FA-sliding pathway, we analysed the protonophoric activity of the corresponding mutants ANT1-R79S, ANT1-R137S, and ANT1-R279S, in which the arginines involved in the substrate-binding site of ANT1 were substituted by serins. ADP/ATP exchange was significantly inhibited only in the ANT1-R79S (Fig. 2b, Suppl. Fig. 1 and Suppl. Table 1), confirming the importance of this amino acid in nucleotide exchange ^23^. Similarly, AA^-^ translocation by ANT1-R79S was by 30 % lower than that of ANT1, whereas the R137S and R279S mutations had little effect on the membrane conductance (Fig. 2c and Suppl. Fig. 2). The addition of 2 mM ATP did not inhibit AA^-^ translocation in ANT-R79S (Fig. 2d and Suppl. Fig. 2), again showing the essential role of R79 for ATP binding. The inhibition was less strong for AN1-K22S, ANT1-R137S and ANT1-R279S as they are also involved in binding of ATP. Based on these data, we conclude that R79 is a second crucial binding site for AA^-^ sliding along the ANT1/membrane interface.

We also tested whether the AA^-^ binds to the protein on the cytosolic side, as suggested by patch experiments in isolated mitoplasts ^7^. The lysine triplet on helix II (K91, K93, and K95) or R79 might represent putative binding sites. For the MD simulation, the AA^-^ was placed in the vicinity of the ANT1 cavity on the cytosolic side. Figure 2e and Video 2 show a characteristic simulation in which the AA^-^ immediately moves away from R79, temporarily binds to the surrounding lysines after approximately 5 ns and ultimately leaves the protein interior after approx. 20 ns. AA^-^ stays outside the protein for the next 80 ns. However, it is possible that AA^-^ remains in the cavity for a longer time due to the presence of positively charged lysines and arginines, but it will eventually exit the cavity because of the unfavorable hydrophobic interactions of the long AA^-^ aliphatic chain and hydrated protein interior.

To test this possibility experimentally, we mutated K93, which points to the lipids, and K22, which is in the protein interior ^16^. Mutation of both lysines gave similar results, with only slight alterations in ADP/ATP exchange (Fig. 2b, Suppl. Fig. 1 and Suppl. Table 1), AA^-^ transport (Fig. 2c and Suppl. Fig. 2) and inhibition by 2 mM ATP (Fig. 2d and Suppl. Fig. 2). Thus, the finding suggests that the two residues don’t form a part of the translocation pathway, in line with MD simulations.

### 3. The AA^-^ is protonated in the hydrated cavity of ANT1

Next, we aimed to identify the final step of AA^-^ sliding to the cytosolic side. Surprisingly, MD simulations revealed no dissociation of the AA^-^ from R79 after 50 ns (Fig. 3a and Video 3). To confirm this, we performed hundreds of independent and unbiased MD simulations (in total over 2 μs), showing that AA was either tightly bound to R79 via a salt bridge or remained in its immediate vicinity. The resulting number density map (Fig. 3b) shows a high probability that AA anion stays in the vicinity of R79. In contrast, if AA was protonated, it readily detached and moved into the lipid bilayer (Fig. 3c and Video 4).

**Figure 3.**
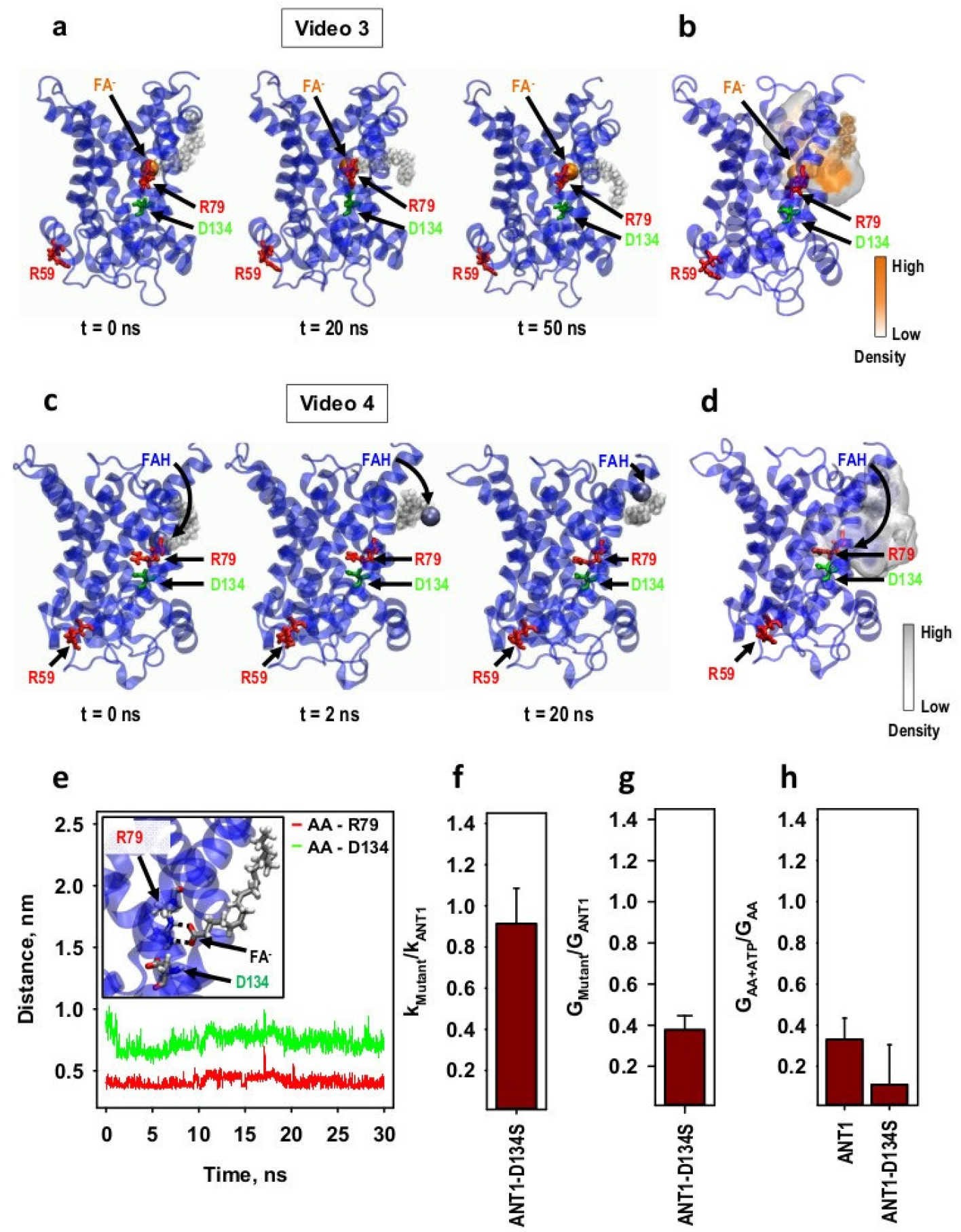
The FA anion (FA^-^) is protonated assisted by D134 while bound to R79. (a) Series of snapshots of the FA^-^ bound to R79 of ANT1, extracted from MD simulations shown in Video 2. Images were taken in the beginning (t = 0 s), after t = 20 ns and at the end of the simulation (t = 50 ns). The FA^-^ tail is shown by van-der-Waals surfaces in grey, the FA^-^ head as a ball in orange. Amino acids R59 (red), R79 (red) and D134 (green) are shown in licorice in the structure of ANT1 (PDB:1OKC). (b) Number density map of the FA^-^ at R79 of ANT1 determined by non-equilibrium MD simulations. (c) Series of snapshots of the protonated FA (FAH) bound to R79, extracted from MD simulations shown in Video 3. Images were taken in the beginning (t = 0 s), after t = 2 ns and at the end of the simulation (t = 20 ns). The labels are similar to (a). Protonated FA head (FAH) is shown as a ball in blue. (d) Number density map of FAH on the protein-lipid interface of ANT1 determined by non-equilibrium MD simulations. (e) Distance between the FA^-^ head and the respective amino acid residues R79 (red) and D134 (green) extracted from the MD simulations in (b). Insert: Salt bridge network of the FA^-^ with R79 and D134 in the vicinity. (f) The ratio of the ADP/ATP exchange rate, k_Mutant_, of the ANT1-D134S to the k_ANT1_ of ANT1, measured with ^3^H-ATP. (g) The ratio of the total membrane conductance, G_Mutant_, of ANT1-D134S to the G_ANT1_ of ANT1, measured in the presence of arachidonic acid (AA) in the membrane. (h) The ratio of the total membrane conductance, G_AA+ATP_, in the presence of AA and ATP to the G_AA_ in the presence of AA only, calculated for the bilayer membranes reconstituted with either ANT1 or ANT1-D134S. Experimental conditions are the same as in Figure 1.

Protonation of AA^-^ in the hydrated cavity of ANT1 thus seems to represent another critical step in the FA-sliding mechanism. The literature shows that an arrangement of positively and negatively charged amino acids promotes the attraction of a proton. The event is crucial for substrate/H^+^ co-transport in several mitochondrial carriers, such as phosphate and aspartate/glutamate carriers ^24-26^. The negatively charged aspartic acid 134 (D134) close to the AA^-^ and R79 (Fig. 3d, e) could facilitate AA^-^ protonation after binding to R79. To test this idea, we substituted D134 by S134 by site-directed mutagenesis. The mutation did not alter ANT1-mediated ADP/ATP exchange (Fig. 3f, Suppl. Fig. 1 and Suppl. Table 1), but AA^-^ transport was reduced by 60 % (Fig. 3g and Suppl. Fig. 2). The inhibition of ANT1-D134S by 2 mM ATP was stronger than the inhibition of ANT1 (Fig. 3g and Suppl. Fig. 2). The effect can be explained by the reduction of one net negative charge in the substrate-binding site of ANT1, which facilitates ATP binding. Our results strongly suggest that protonation of the AA^-^ is possible with the support of D134, which is located close by.

### 4. R79 mediates the competition of AA^-^ and substrate transport in ANT1

R79 is important for the binding of AA^-^ as well as for the substrates ATP and ADP (Fig. 4a) and the inhibitors CATR (Fig. 4b) and BKA (Fig. 4c) of ANT1. We further investigated whether R79 mediates the competition between the AA^-^ and ATP, CATR and BKA. Fig. 4d and Suppl. Fig. 3a show that ADP/ATP exchange is negatively regulated by the presence of AA in a concentration-dependent manner. The ADP/ATP exchange of ANT1-R79S was generally lower but unaffected by the presence of AA (Fig. 4d and Suppl. Fig. 3b). The addition of 100 μM ATP changed the membrane conductance for ANT1 but not ANT1-R79S, with the effect depending on the AA concentration in the membrane (Fig. 4e and Suppl. Fig. 4). Similarly, the AA content had a strong effect on the inhibition of ANT1 by 10 μM CATR and BKA but far less effect on the inhibition of ANT1-R79S (Fig. 4f, Fig. 4g and Suppl. Fig. 4). Thus, ANT-mediated ADP/ATP exchange and AA^-^ transport are interdependent and negatively regulate each other by competitive binding at R79.

**Figure 4.**
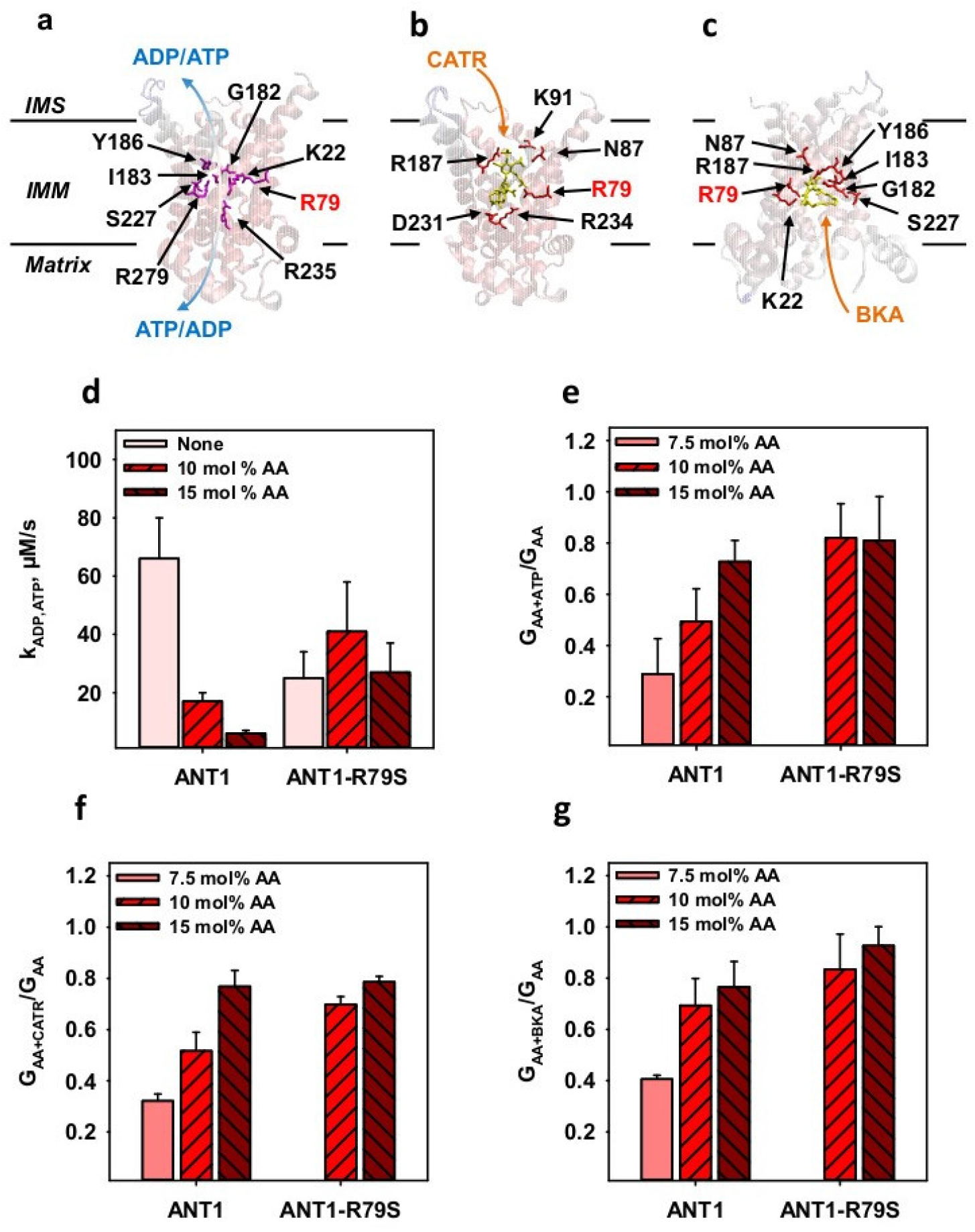
Competition between the fatty acid anion (FA^-^) and the ANT-specific substrates is mediated by the binding to R79 in ANT1. Crystallographic structures of ANT1 in complex with carboxyatractyloside (CATR; PDB:1OKC) (a, b) or bongkrekic acid (BKA; PDB:6GCI) (c) are shown to indicate the localization of amino acids involved in the binding of ATP (a), CATR (b) or BKA (c). Abbreviations are similar to Figure 1. (d) ADP/ATP exchange rates (k_ADP, ATP_) of ANT1 or ANT1-R79S in the presence (10 and 15 mol%) and absence of the arachidonic acid (AA) in the membrane measured with ^3^H-ATP. (e - g) The ratio of the total membrane conductance of the membranes reconstituted with ANT1 or ANT1-R79S in the presence of AA and 100 μM ATP (G_AA+ATP,_ (e)), or 10 μM CATR (G_AA+CATR,_ (f)) or 10 μM BKA (G_AA+BKA,_ (g)) to the total membrane conductance in the presence of AA only (G_AA_). In all experiments, the lipid concentration was 4 mg/ml (d) or 1.5 mg/ml (e - g) and the protein concentration was 4 μg/mg of lipid (d - g). Membranes were made of PC:PE:CL (45:45:10 mol%) and reconstituted with AA in concentrations indicated in the figures. Buffer solution contained 50 mM Na_2_SO_4_, 10 mM Tris, 10 mM MES and 0.6 mM EGTA at pH = 7.34 and T = 296 K (d) or T = 306 K (e - g). Data are the mean ± SD of at least three independent experiments.

## Discussion

We report the molecular mechanism by which ANT1 transports FA^-^ across the inner mitochondrial membrane, in accordance with the FA cycling model. After the initial

“trapping” by the positively charged region around R59, the FA^-^ translocates along the protein-lipid interface such that the negatively charged FA carboxylic group is in close contact with the positively charged surface, while the hydrophobic chain is shifted through the hydrocarbon layer of the lipid membrane adjacent to the protein. The effect is similar to the “credit card” model proposed for lipid scramblases ^27,28^. Thus, ANT1 is a carrier for FA^-^ that fits the established view that all functionally characterized members of the mitochondrial carrier family SLC25 are anion carriers ^29,30^.

According to the FA cycling hypothesis, FA binds on the protein’s matrix side (m-side). The similarity of the H^+^ transport characteristics between ANT1 and UCPs implies that the FA^-^ may be trapped by positively charged amino acids on the m-side of ANT1, as proposed for UCP2 ^31,32^. The FA^-^ initially binds at R59 as a crucial part of the suggested “sliding pathway” (Fig. 5, (1)). R59 is the homolog of R60 in UCP2, to which FA^-^s were shown to bind ^31,32^. Our results contradict previous reports that FAs can only bind from the cytosolic side (c-side) of the protein ^33^. Although in principle possible, FA binding seems unnecessary as the FA^-^ can easily be protonated on the cytosolic side and its neutral form can rapidly diffuse through the inner mitochondrial membrane ^12,13^. The initial binding on the m-side starts the FA cycling and, thus, mitochondrial uncoupling to counteract the generation of reactive oxygen species. The transport of FA anions to the cytosol is of great physiological significance as it prevents the accumulation of free FAs in the mitochondrial matrix. Excessive amounts of FAs induce lipotoxicity and are involved in the pathogenesis of obesity, diabetes, ischemia, and degenerative diseases ^34-38^.

**Figure 5.**
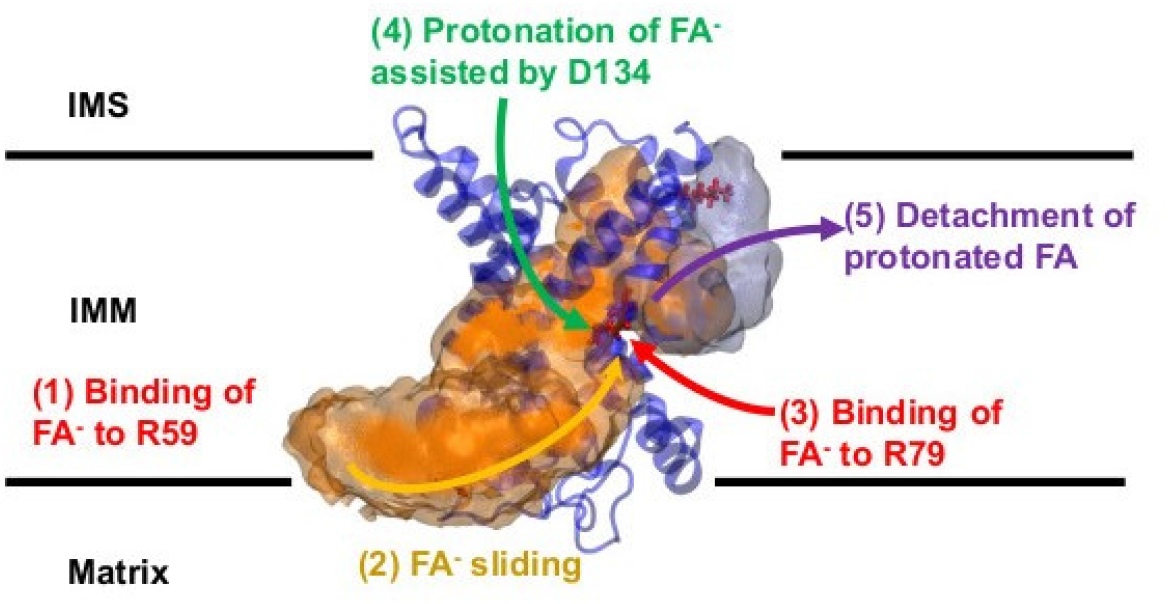
The mechanism of FA anion (FA^-^) transport by ANT1. The FA anion initially binds to R59 of ANT1 from the matrix side (1). From this binding site, it slides along the positively charged protein-lipid interface to R79 inside the inner mitochondrial membrane (IMM) (2). The FA anion binds to R79 (3), where it receives a proton from the hydrated cavity assisted by D134 (4). The protonated FA finally detaches from the protein and moves into the membrane to the intermembrane space (IMS) (5).

After FA^-^ binds to the matrix side, it follows the positively charged surface of ANT1 (Fig. 5, (2)) as proposed earlier ^6^. Longer and more unsaturated FA diffuse faster through the lipid bilayer and their p*K*a in lipid bilayers is closer to the physiological pH ^39^. pH is crucial to accelerate protonation and deprotonation at the lipid-water interface ^40^. The presence of a hydrophobic pocket in the protein, in which longer and more unsaturated FA would reside longer, would lead to the same finding but the resolved structures of ANT1 show no such hydrophobic pocket, ruling out this possibility ^16,17^. Thus, it is far more likely that the FA anion translocates along the protein-lipid interface.

Several membrane proteins (flippases and scramblases) are already known to mediate the flip-flop of phospholipids or FAs across lipid bilayers by the “credit-card” mechanism ^41-45^. This is based on the crystallographic structure of the calcium-activated chloride channel (CaCC/TMEM16) and the lysosomal integral membrane protein 2 (LIMP2), which function as a lipid scramblase or a FA translocase ^28,46,47^. Both proteins contain a membrane-spanning hydrophilic groove at the protein-lipid interface, in which the phospholipid/FA head is inserted and transported along, while the hydrophobic tail remains inside the lipid bilayer. However, lipid transfer through the groove is coupled to the leakage of lipid-conjugated ions such as Ca^2+^, making a similar mechanism unsuitable for transporting FA^-^ across the inner mitochondrial membrane. Work with modified lipid heads of various sizes has suggested that lipid scrambling also occurs on the protein-lipid interface outside the groove of CaCC, which excludes the transport of lipid-conjugated ions ^27,48^. Similarly, our MD simulations show that the FA^-^ moves along the lipid-protein interface from R59 to R79 independent of a hydrophilic grove, driven mainly by the electrostatic surface potential of ANT1. In this way, FA^-^ transport is not associated with any unwanted ion leakage across the IMM and mitochondrial function is not compromised.

We have found a second FA^-^ binding site - R79 - in the substrate-binding site of ANT1 (Fig. 5, (3); ^49,50^. This is of particular interest because the substrates (ADP and ATP) and the inhibitors (CATR and BKA) use R79 as a common binding target ^16,18,19,51-53^. Not only does ADP/ATP exchange negatively regulate FA transport ^7^, FA^-^ and the substrates (CATR and BKA) compete for R79. In the presence of FA, ADP/ATP exchange is diminished for ANT1 but not for ANT1-R79S. We conclude that FA does not displace the ADP or ATP^-^ but decreases the binding affinity for these substrates, as the R79 residue is blocked for as long as the FA^-^ is bound. Another group has postulated the existence of a similar binding target in the central cavity of the carriers UCP1 and ANT1 but the precise amino acid has remained unknown ^14,15,33^. The role of the FA^-^ is different in the two models. (i) FAs are co-factors that bind to ANT1 or UCP1 to enhance H^+^ translocation through the centre of the proteins. Both protonation and deprotonation of the FA are thought to occur inside the protein. In contrast, (ii) the FA cycling mechanism holds that protonation/deprotonation of the FA only occur at the lipid-water interface ^11^.

We have found strong evidence for a novel and critical third step in the FA sliding model. Instead of being further transported to the cytosolic side, the FA^-^ is protonated while bound to R79 (Fig. 5, (4)). R79 attracts the FA^-^ carboxylic head into the hydrated protein cavity and the spatially close D134 mediates proton transfer from the hydrated cavity to the FA^-^. Several mitochondrial carriers that co-transport H^+^ and substrates, such as the phosphate carrier ^25^, the aspartate/glutamate carrier ^26^, the oxodicarboxylate carrier ^54^ and the GTP/GDP carrier could use a similar protonation/deprotonation mechanism ^55^. Small amounts of H^+^ are moved together with the ADP/ATP exchange, although proton exchange does not invariably occur ^56,57^. In the final step, the neutral FAs are immediately released into the membrane (Fig. 5, (5)) and the catalytic cycle can start again.

The FA^-^ translocation mechanism of ANT1 is energetically more favourable than the FA co-factor or shuttling model proposed for ANT1 and UCP1 ^15,33^. Our MD simulations show that long-chain FAs do not spontaneously bind to R79 when initially placed on the c-side and entering the hydrated ANT1 cavity. In contrast, the configuration is stable when FAs bind from the m-side to R79. Furthermore, if the FA promotes H^+^ transport through the protein centre by either FA flip-flop or as a co-factor, the FA carboxylic group carrying an H^+^ has to cross at least one salt bridge network that closes ANT1 either to the cytosolic or to the matrix side ^58^. As a part of the alternating access mechanism ^17^, salt bridge networks must function properly and ANT1 does not undergo a spontaneous conformational change in the absence of ADP or ATP ^20,59^, which is assumed to require around 10 kcal mol^-1 60^. The energy released by the binding of the singly charged FA^-^ or H^+^ is too low to break the salt bridge network, in contrast to the large amount of energy released upon binding the multiply negatively charged ATP and ADP anions to the positively charged bottom of the protein cavity. In addition, MD simulations of ANT1 and UCP2 in DOPC bilayers have shown that water, including H_3_O^+^, is hindered by the cytosolic or matrix salt bridge network and cannot leak through the protein ^20,32^. A substantially different model proposes that UCP2 forms tetramers, mediating FA-activated H^+^ transport ^61^. Although not investigated in detail, ANT1 tends to form dimers^62^. In contrast, we show that FA can be transported through the ANT1 monomer and oligomerization is not a prerequisite for transport, although we cannot exclude the possibility that it favours FA^-^ transport over substrate transport.

The ANT1-mediated FA^-^ transport we describe involves amino acids that are well-conserved among SLC25 members ^63^. We suggest that the mechanism of FA^-^ transport also applies to other mitochondrial carriers that show protonophoric activity in the presence of FA, such as the uncoupling proteins, the dicarboxylate carrier and the aspartate/glutamate carrier ^64-66^. Indeed, the activation and inhibition pattern of UCP1-3 also points to the FA cycling model ^3,9,10^. The precise mechanism of FA^-^ transport might differ and the mechanistic details of each mitochondrial carrier await elucidation.

## Materials and methods

### Chemicals

Sodium sulfate (Na_2_SO_4_), 2-(N-morpholino)ethanesulfonic acid (MES), tris(hydroxymethyl)-aminomethane (Tris), sodium dodecyl sulfate (SDS) ethylene glycol-bis(β-aminoethyl ether)-N,N,N’,N’-tetraacetic acid (EGTA) were purchased from Carl Roth GmbH & Co. K.G. (Karlsruhe, Germany). Hexane, hexadecane, dimethyl sulfoxide (DMSO), purine nucleotides adenosine tri-, di-, and guanosine triphosphate (ATP, ADP, GTP), agarose, carboxyatractyloside (CATR), and bongkrekic acid (BKA) were purchased from Sigma-Aldrich Co. (Vienna, Austria). 1,2-dioleoyl-sn-glycero-3-phosphocholine (DOPC), 1,2-dioleoyl-sn-glycero-3-phosphoethanolamine (DOPE) and cardiolipin (CL) came from Avanti Polar Lipids Inc (Alabaster AL, USA) and chloroform from Sanova Pharma GesmbH (Austria). Arachidonic acid (AA) was purchased from Larodan (Biozol, Eching, Germany).

### Cloning, site-directed mutagenesis, isolation, and reconstitution of murine ANT1

Cloning, isolation and reconstitution of murine ANT1 and ANT1 mutants followed an established protocol ^67^. In brief, for protein expression we used E. coli strain Rosetta (DE3; Novagen). To isolate inclusion bodies we centrifuged cells disrupted by high-pressure cell disrupter *One Shot* (Constant Systems Limited, Daventry, UK) at 1 kbar. For reconstitution, protein from inclusion bodies was first solubilized in 100 mM Tris at pH 7.5, 5 mM EDTA, 10% glycerin (TE/G-buffer) containing 2% sodium lauryl sulfate and 1 mM DTT. Then, it was gradually mixed with the membrane-forming lipids (DOPC, DOPE and CL; 45:45:10 mol%) dissolved in TE/G-buffer containing 1.3% Triton X-114, 0.3% n-octylpolyoxyethylene, 1 mM DTT and 2 mM GTP. After several dyalisis steps the mixture was dialyzed three times against assay buffer (50 mM Na2SO4, 10 mM MES, 10 mM Tris, 0.6 mM EGTA at pH 7.35). To eliminate aggregated and unfolded proteins, the dialysate was centrifuged and run through a hydroxyapatite-containing column (Bio-Rad, Munich, Germany). Non-ionic detergents were removed using Bio-Beads SM-2 (Bio-Rad).

To generate ANT1 mutants, we carried out site-directed mutagenesis of lysine 48, 51, 62, 93, arginine 59, 79, 137, 279, and aspartic acid 134 to serine using the Q5 Site-Directed Mutagenesis Kit (New England Biolabs, Austria). The sequences were verified by both DNA and amino acid sequencing. Mutants were always refolded in parallel with a wildtypes to minimize artefacts. Critical mutants such as R59S and R79S were refolded at least twice to rule out any possible side-effects of the refolding.

The protein concentration of the proteoliposomes was measured with the Micro BCATM Protein Assay Kit (Thermo Fisher Scientific, Prod. #23235). SDS-PAGE and silver staining verified protein purity. Furthermore, we have performed the control measurements of ADP/ATP exchange, activation by free fatty acids and inhibition by 2 mM ATP of both, the mutant and the parallel refolded wildtype to ensure the protein functionality.

### Exchange rate measurements of ANT1

ANT1-mediated exchange of ATP/ADP was measured radioactively using ^3^H-ATP (Perkin Elmer Prod.# NET420250UC) ^67^. The standard protocol was slightly modified in that ^3^H-ATP was initially present in the proteoliposomes to measure the release of the radionucleotide over time (Supplementary Fig. 1).

### Electrophysiological measurements of ANT1

Planar lipid bilayers were formed at the top of the dispensable plastic pipette as described previously ^68^. AA was added directly to the lipid phase before membrane formation. We verified proper membrane formation by measuring the membrane capacitance (C = 0.72 ± 0.05 μF/cm^2^). C was independent of the presence of protein, FA and inhibitor. Current-voltage (I-U) recordings were performed with a patch-clamp amplifier (EPC 10USB, Harvard Bioscience). Total membrane conductance (G_m_) at 0 mV was derived from the slope of a linear fit of the experimental data at applied voltages from -50 mV to + 50 mV (Supplementary Fig. 2, a). ATP in buffer solution (pH = 7.34) and the ANT-specific inhibitors BKA and CATR (in DMSO) were added to the buffer solution before forming bilayer membranes. The volume of the added inhibitors in DMSO did not exceed 10 μl and did not alter total membrane conductance as previously shown ^6^. The concentrations of each substrate are indicated in the Figures. Membrane conductance expressed in relative units was calculated as previously described ^10^.

### Molecular dynamics simulations

We performed all-atom molecular dynamics (MD) simulations of ANT1 protein in a 1,2-dioleoyl-sn-glycero-3-phosphocholine (DOPC) bilayer with the addition of AA (20:4). Residues 1 and 294-297 missing from the crystal structure of ANT1 (PDB code: 1okc)^16^ without CATR were added using Modeller9 ^69^ and implemented into the DOPC bilayer using CHARMM-GUI (http://www.charmm-gui.org/) ^70-72^. Additionally, three cardiolipin (CL) molecules were introduced in the system to surround the protein at the crystallographically determined positions ^16^. Initial simulation box contained ANT1 protein (with a total charge of +19), 73 DOPC molecules per leaflet (146 per system), ∼11500 water molecules, three CL molecules and the necessary number of Cl anions to neutralize the net charge. The prepared system was first minimized and equilibrated in six steps using the CHARMM-GUI protocol ^73^, and then simulated for a further 100 ns without any restraints with a time step of 2 fs in a periodic rectangular box of 7.9 nm x 7.9 nm x 9.4 nm using the isobaric-isothermal ensemble (NPT) and periodic boundary conditions in all directions at T = 310 K, maintained via a Nosé–Hoover thermostat ^74^ independently for the DOPC, water/ions and protein subsystems with a coupling constant of 1.0 ps^-1^. The pressure was set to 1.013 bar and controlled with a semi-isotropic Parrinello-Rahman barostat ^75^ with a time constant for pressure coupling of 5 ps^-1^. We used the particle-mesh Ewald (PME) method ^76^ to calculate long-range electrostatics. Real-space Coulomb interactions were cut off at 1.2 nm using a Fourier spacing of 0.12 nm and a Verlet cut-off scheme.

Analysis of an electrostatic potential map of ANT1 revealed a tentative translocation pathway of charged AA species, namely the path following positive electrostatic potential (see Figure 1b) corresponding to regions abundant in arginine and lysine residues. We set up 18 initial configurations of the ANT1 – AA system, where AA^-^ molecules were placed in 18 distinct positions roughly along the tentative pathway, encompassing around 1/5 of the entire surface area of the ANT1 protein. We propagated 20 independent and unbiased runs for each of the 18 initial configurations. Each run started with randomly generated velocities of atoms that were obtained from the Boltzmann distribution at 310 K and propagated for 20 ns. Combined data from all independent unbiased simulations (18 sets × 20 runs/set × 20 ns gives an overall sampling time of 7.2 μs) enabled us to generate a 3D number density distribution of carbon atom from a carboxylic group of AA^-^ surrounding ANT1. Density distribution was obtained using the VMD molecular graphics program, with average density distribution of all snapshots belonging to all 360 (18 sets × 20 runs/set) propagated simulations shown in Figure 2a (density plot obtained using isovalue = 30). Upon finding a hotspot, namely the region around the R79 residue, we propagated 50 ns long simulations starting with AA^-^ bound to R79 (Figure 3a). Moreover, we used four configurations of the ANT1 – AA^-^ system found in the previous steps to monitor the behavior of the neutral form of AA in the same region, i.e. around the R79 hotspot. We propagated 25 independent runs for each of the four configurations of the ANT1 – AA system for 20 ns (atom velocities were again randomly generated at the initialization of the simulations), giving rise to the 3D number density distribution of the carbon atom in the carboxyl group of the AA shown in Figure 3c and Figure 5 (blue region). We prepared force field parameters (CHARMM36m) to describe neutral AA by adding hydrogen atom to the carboxylic group of the AA^-^ molecule in the four chosen starting configurations (one chloride atom was added to the system in these simulations to preserve charge neutrality). Finally, to test whether AA^-^ binds to R79 from inside of the protein cavity, we set up 100 ns long simulation starting with AA^-^ bound via its carboxylic group to R79 (Figure 2e).

All simulated systems were described by the CHARMM36m force field ^77^. The electrostatic potential maps of all systems were calculated with the PMEPOT plugin of VMD ^78^. All simulations were run with the GROMACS 5.1.4 software package ^79^ and visualized with the VMD molecular graphics program ^80^.

### Statistics

Data from the electrophysiological measurements are displayed as mean ± S.D. of at least three independent experiments. Each experimentally derived result was the mean membrane conductance from a minimum of two formed bilayer membranes. The recombinant proteins from at least two refoldings were usually used to ensure the reproducibility of the experiments.

## Data availability

The data supporting the findings are available within the paper, in the Supplementary Information and from the corresponding author upon reasonable request.

## Acknowledgments

We are grateful to Peter Pohl (Institute of Biophysics, University of Linz, Austria) and Yury N. Antonenko (A. N. Belozersky Institute of Physico-Chemical Biology, Moscow State University, Russia) for valuable discussions, and to Dr Graham Tebb for editorial assistance. We thank the computer cluster Isabella based in SRCE - the University of Zagreb, University Computing Centre for computational resources. Finally, we thank Tatiana Tyshchuk and Max Bauernfeind for their excellent technical assistance.

## Authors contributions

J.K., M.V., and E.E.P. designed the study, S.B. produced and validated the mutants, J.K. undertook the electrophysiological experiments, Z.B., S.Š. and M.V. performed molecular dynamics simulations, J.K., S.Š., Z.B., and M.V. analysed the data and prepared the figures, J.K., M.V. and E.E.P. interpreted the data and wrote the manuscript. All authors revised and agreed on the final version.

