## Supplementary Figures for "Mechanism of the ANT-mediated transport of fatty acid anions across the inner mitochondrial membrane"

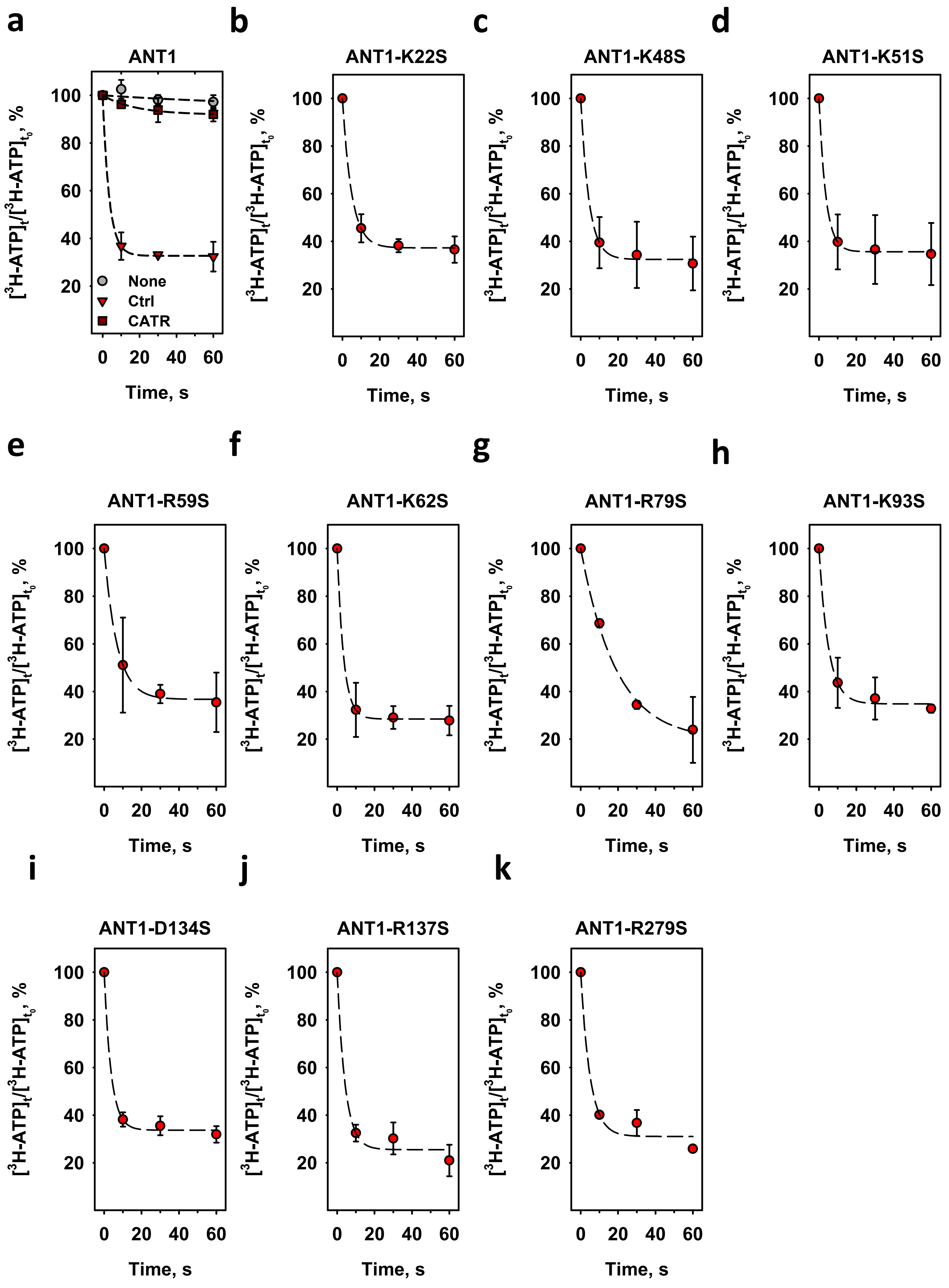


**Supplementary Figure 1.**

**Supplementary Figure 1. Release of $^3$H-ATP from liposomes reconstituted with ANT1 or mutants.**

Time course of the $^3$H-ATP concentration in liposomes in the absence (grey circles) and in the presence of reconstituted ANT1 (red) (a), ANT1-K22S (b), ANT1-K48S (c), ANT1-K51S (d), ANT1-R59S (e), ANT1-K62S (f), ANT1-K93S (g), ANT1-R79S (h), ANT1-D134S (i), ANT1-R137S (j) or ANT1-R279S (k). Concentration of CATR in (a) was 100 µM (dark red). Lines represent a least square regression fit of an exponential function to the data. In all experiments, the lipid concentration was 1.5 mg/ml and the protein concentration was 8.5 - 9 µg/mg of lipid. Membranes were made of PC:PE:CL (45:45:10 mol%). Buffer contained 50 mM $Na_2SO_4$, 10 mM Tris, 10 mM MES and 0.6 mM EGTA at pH = 7.34 and T = 296 K. ATP and ADP were dissolved in buffer solution and adjusted to pH = 7.34. Data are shown as the mean ± SD from at least three independent experiments.

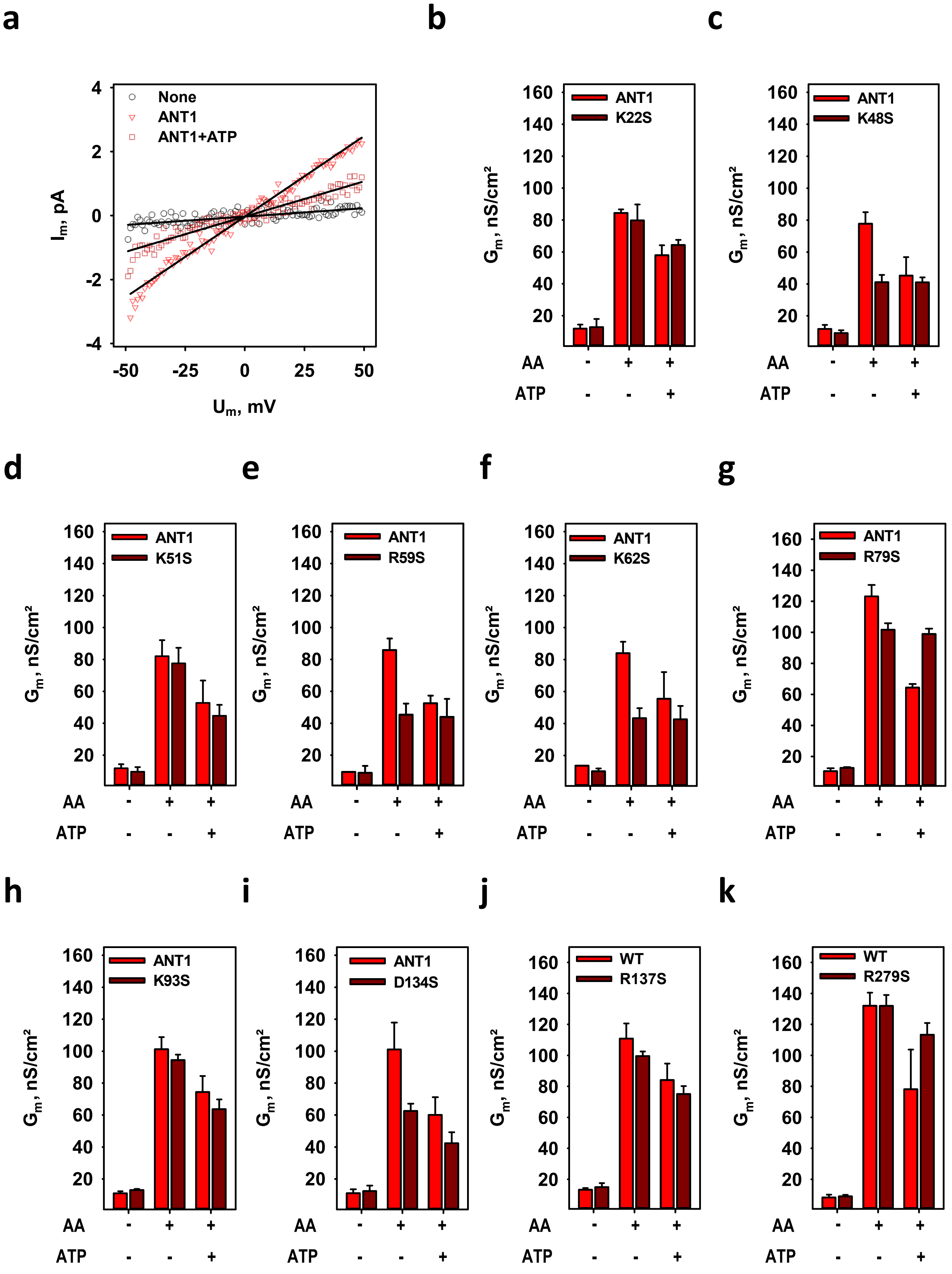


**Supplementary Figure 2**

**Supplementary Figure 2. Electrical parameters of planar lipid bilayers reconstituted with ANT1 and mutants.**

(a) Representative current-voltage recording of lipid bilayer membranes reconstituted with ANT1 in the absence of arachidonic acid (AA, black circles), in the presence of AA (red triangles) and in the presence of AA and 2 mM ATP (dark red squares). Total membrane conductance ($G_m$) is the slope of a linear fit to the data. (b - k) $G_m$ of the planar lipid bilayers reconstituted with the ANT1-K22S (b), ANT1-K48S (c), ANT1-K51S (d), ANT1-R59S (e), ANT1-K62S (f), ANT1-R79S (g), ANT1-K93S (h), ANT1-D134S (i), ANT1-R137S (j) and ANT1-R279 (k) and parallel refolded ANT1 in the absence of AA or in the presence of AA or in the presence of AA and 2 mM ATP (third data set). In all measurements, lipid concentration was 1.5 mg/ml and the protein concentration - 4 µg/(mg of lipid). Membranes were made of PC:PE:CL (45:45:10 mol%) reconstituted with AA in concentration indicated in the figures. Buffer contained 50 mM $Na_2SO_4$, 10 mM Tris, 10 mM MES and 0.6 mM EGTA at pH = 7.34 and T = 306 K. ATP was dissolved in buffer solution and adjusted to pH = 7.34. Data are displayed as the mean ± SD of at least three independent measurements.

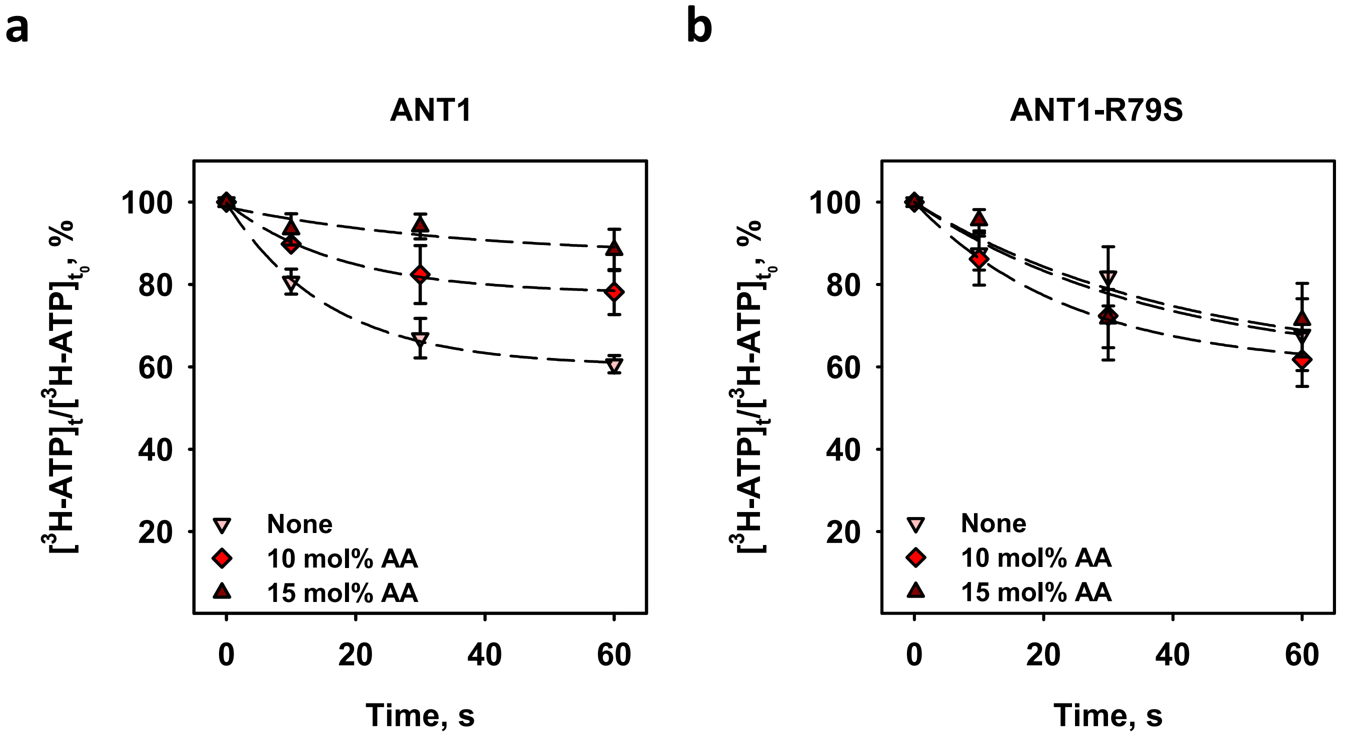


**Supplementary Figure 3. Release of $^{3}$H-ATP from liposomes reconstituted with ANT1 or ANT1-R79S.**

Time course of the relative $^{3}$H-ATP concentration measured in proteoliposomes reconstituted with ANT1 (a) or ANT1-R79S (b) in the absence (none) or in the presence of 10 or 15 mol% AA. Lines are a least-square regression fit of an exponential function to the data.

In all experiments, the lipid concentration was 4 mg/ml and the protein concentration was 4 µg/(mg of lipid). Membranes were made of PC:PE:CL (45:45:10 mol%) reconstituted with AA as indicated in the figures. Buffer solution contained 50 mM $Na_2SO_4$, 10 mM Tris, 10 mM MES and 0.6 mM EGTA at pH = 7.34 and T = 296 K. Data are the mean ± SD of at least three independent experiments.

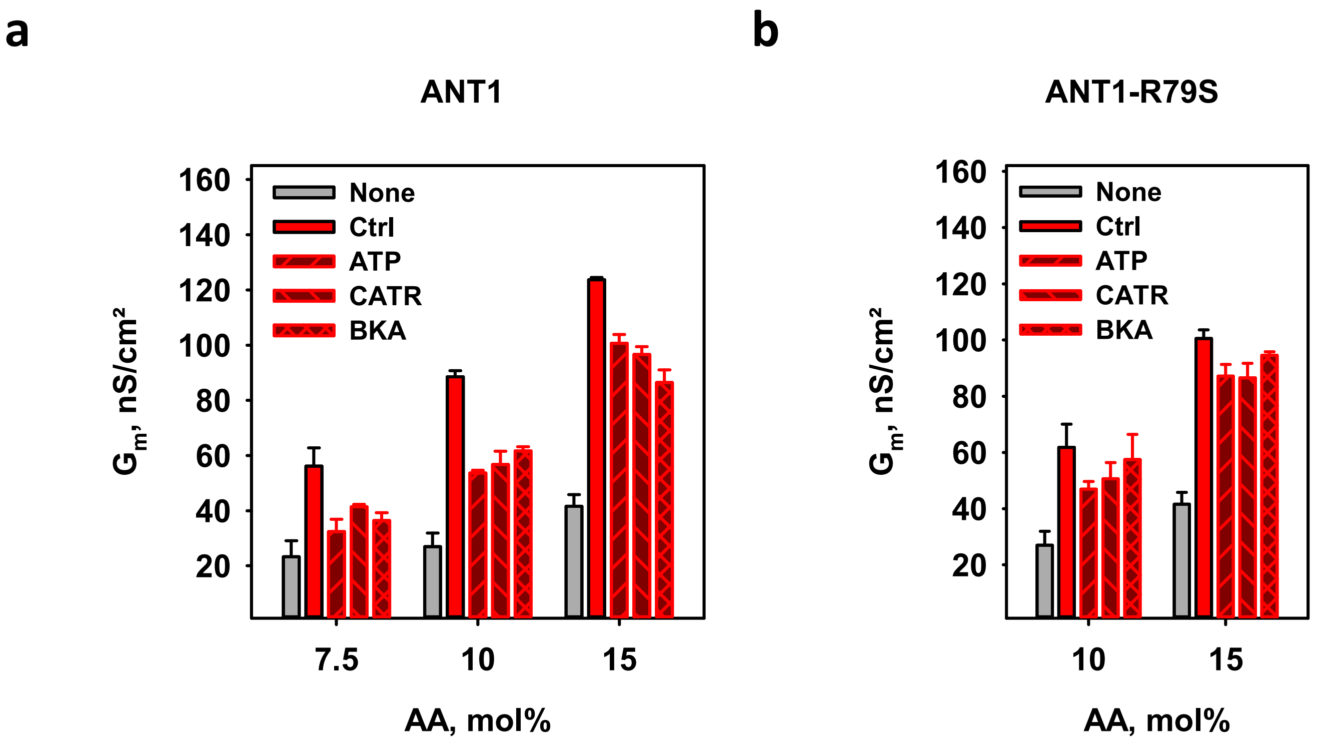


**Supplementary Figure 4. Total membrane conductance of planar lipid bilayers reconstituted with ANT1 or ANT1 R79S**

Total membrane conductance ($G_m$) of planar lipid bilayers reconstituted with ANT1 (a) or ANT1-R79S (b) in the presence of 7.5, 10 or 15 mol% AA. Two control experiments were made in the absence of ANT1 and inhibitors (none) or in the presence of ANT1 only (ctrl). The inhibitors were added in following concentrations: 100 µM ATP, 10 µM CATR or 10 µM BKA. In all experiments, the lipid concentration was 1.5 mg/ml and the protein concentration was 4 µg/mg of lipid. Membranes were made of PC:PE:CL (45:45:10 mol%) reconstituted with AA as indicated in the figures. Buffer solution contained 50 mM $Na_2SO_4$, 10 mM Tris, 10 mM MES and 0.6 mM EGTA at pH = 7.34 and T = 306 K. ATP was dissolved in buffer and pH adjusted to 7.34, CATR and BKA were dissolved in DMSO. Data are the mean ± SD of at least three independent experiments.

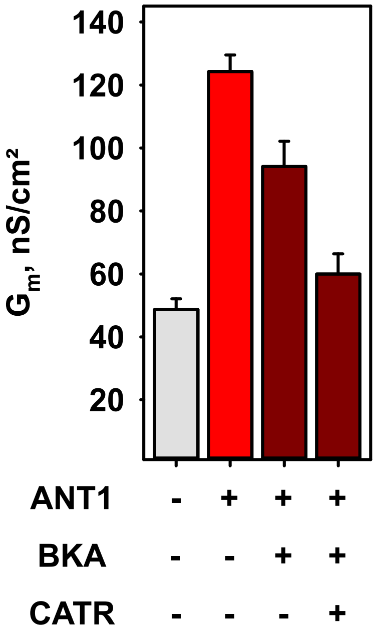


**Supplementary Figure 5. Inhibition of ANT1-mediated FA anion transport by specific inhibitors of ANT1.**

Total membrane conductance ($G_m$) of planar lipid bilayers in the presence of 15 mol% arachidonic acid (AA). BKA and CATR were added in concentration of 100 µM. In all experiments, the lipid concentration was 1.5 mg/ml and the protein concentration was 4 µg/mg of lipid. Membranes were made of PC:PE:CL (45:45:10 mol%), reconstituted with AA. Buffer solution contained 50 mM $Na_2SO_4$, 10 mM Tris, 10 mM MES and 0.6 mM EGTA at pH = 7.34 and T = 306 K. BKA and CATR were dissolved in DMSO and subsequently added to the bulk solution. Data are the mean ± SD of at least three independent experiments.
